# Characterization of five environmental phages infecting *Escherichia coli* K-12 isolated during a phage biology training course

**DOI:** 10.1101/2025.08.13.670052

**Authors:** Lena Shyrokova, Artyom A. Egorov, Astrid Cole, Julian J. Duque-Pedraza, Anu Tyagi, Karin Ernits, Toomas Mets, Tatsuaki Kurata, Jonas Juozapaitis, Alessio Ling Jie Yang, Rodrigo Ibarra Chavez, Frederik Oskar Graversgaard Henriksen, Swenja Lassen, Gemma C. Atkinson, Vasili Hauryliuk, Marcus J.O. Johansson

**Affiliations:** Department of Experimental Medical Science, Lund University, Lund, Sweden; NanoLund, Lund University, Lund, Sweden; Institute of Technology, University of Tartu, Tartu, Estonia; Science for Life Laboratory, Lund, Sweden; Cryo-EM for Life Sciences, Lund University, Lund, Sweden; RNA Systems Biochemistry Laboratory, Pioneering Research Institute, RIKEN, 2-1 Hirosawa, Wako, Saitama 351-0198, Japan; UAB Seqvision, Budiniškių g. 3-9, LT-05275 Vilnius, Lithuania; European Molecular Biology Laboratory, Genome Biology Unit, Heidelberg, Germany; Collaboration for joint PhD degree between EMBL and Heidelberg University, Faculty of Biosciences, Heidelberg, Germany; Section of Microbiology, University of Copenhagen, Copenhagen, Denmark; Department of Molecular Biology and Genetics, Aarhus University, Aarhus, Denmark; Research Group for Food Microbiology and Hygiene, National Food Institute, Technical University of Denmark, Kongens Lyngby, Denmark

## Abstract

Phage collections are essential tools for discovering and dissecting bacterial anti-phage defense systems. Here, we report the isolation and characterization of five environmental *Escherichia coli*-infecting phages, obtained during the 2023 *Fundamentals of Basic and Applied Phage Biology* course at Lund University. The phages were isolated using a motile *E. coli* K-12 BW25113 strain, whose motility is conferred by an IS5 insertion upstream of the *flhDC* operon, the master regulator of flagellar synthesis. The isolated *Escherichia* phages include Lubas (LuPh1) and Lucat (LuPh2) of the genus *Tequatrovirus*; Lupin (LuPh3) and Lucris (LuPh4) of the genus *Tequintavirus*; and Kompetensportalen (LuPh5) of the genus *Chivirus*. Transmission electron microscopy confirmed myovirus and siphovirus morphologies consistent with these genera. As expected for phages in the flagellotropic *Chivirus* genus, LuPh5 failed to infect a poorly motile BW25113 strain lacking the IS5 element upstream of *flhDC*. By testing a panel of eight previously described anti-phage defense systems, we found that LuPh1 and LuPh2 are inhibited by the toxin-antitoxin-chaperone CmdTAC system; LuPh5 is inhibited by both the restriction-modification system EcoRI and the abortive infection reverse transcriptase AbiK; and all five phages are sensitive to the hybrid artificial CmdTA-HigC system. Collectively, our findings expand the toolkit for probing phage-host interactions and underscore the pedagogical value of incorporating phage isolation into practical training for emerging researchers.

## Introduction

Bacteriophages, or simply phages, are the viruses of bacteria. To defend against viral attacks, bacteria employ diverse defense systems (1–3). Phages, in turn, can counter these defenses with specialized anti-defense mechanisms (4). Phage collections assembled by numerous labs are an important resource for identification and characterization of defense and anti-defense systems (5–8). An example of a freely shared and broadly used collection is the BActeriophage SElection for your Laboratory (BASEL), a set of *Escherichia coli*-infecting phages isolated by the Alexander Harms lab (9, 10). Finally, phages are also available via repositories, such as at the Leibniz Institute DSMZ and at the Félix d’Hérelle Reference Center for bacterial viruses at the Université Laval.

The isolation of novel phages is, at least for some bacterial species, relatively straightforward, making it well-suited for course-based research experiences (CREs). The most famous ‘phage hunt’ CRE is organized by The Science Education Alliance (SEA). The SEA-PHAGES (SEA Phage Hunters Advancing Genomic and Evolutionary Science) is a large-scale operation (11). Over the years, more than 50,000 undergraduate researchers have isolated 23,000 bacteriophages for 16 different genera of Actinobacteria, including *Mycobacterium smegmatis*, *Gordonia terrae* and *Microbacterium foliorum*, and over 4,500 of these phages have been fully sequenced and annotated. Phages from the collection were successfully used for phage therapy against *Mycobacterium* infections (12). Even small-scale phage hunting CREs can be impactful. An example is the Durham collection of 12 *E. coli*-infecting phages that were isolated during undergraduate practical classes and used as tools to provide mechanistic insights into the Bacteriophage Exclusion (BREX) anti-phage defense systems (13, 14).

In April 2023, our lab at Lund University organized the *Fundamentals of Basic and Applied Phage Biology* course (April 24–28) inspired by the renowned Cold Spring Harbor Phage Course (Susman, 1995). It was designed for PhD students and postdocs and held within the framework of the Swedish National Doctoral Programme in Infection and Antibiotics (NDPIA). The aim of the course was to provide the participants with a comprehensive theoretical overview of both fundamental and applied phage biology, complemented with a set of experimental skills that would allow the introduction of phage techniques into an ongoing research program. Phage experts from across Europe delivered lectures on topics ranging from phage diversity and environmental distribution to their ecological roles, interactions with bacterial hosts, mechanisms of bacterial immunity, and the development of phage-based antibacterial treatments. Laboratory work included the isolation of *E. coli*-infecting phages from the environment, determination of phage titers, anti-phage defense assays, as well as the selection of mutant phages resistant to an anti-phage defense system.

In this study, we characterize five environmental coliphages isolated during the NDPIA course sampling and laboratory sessions. We present their annotated genomes, taxonomic classification, morphological features, lytic activity, and susceptibility to eight previously described anti-phage defense systems.

## Results and Discussion

To increase the chances of successfully isolating *E. coli*-infecting phages from environmental water samples, we used the laboratory *E. coli* K-12 strain BW25113 as the host (15, 16). Laboratory *E. coli* K-12 strains lack O-antigen polysaccharides, which results in exposed surface receptors and enhanced susceptibility to a wide range of phages (17, 18). Moreover, BW25113 carries a mutation in the *hsdR* gene, which inactivates the type I restriction-modification (R-M) system EcoKI (15, 16). It should, however, be noted that BW25113 harbors additional anti-phage defense systems (10), which may prevent the isolation of certain phages. To confirm the identity of the BW25113 strain in our collection (designated VHB17), we sequenced its genome using both Illumina and Oxford Nanopore technologies. The analyses verified the strain’s identity and revealed a few differences compared to the reference BW25113 genome (Table S1). These differences included the presence of an IS5 element in the intergenic region between *uspC* and *flhDC* (Fig. 1A). Although BW25113 is considered a largely non-motile strain, the insertion of IS elements in this region is known to increase motility by elevating expression of the *flhDC* operon, encoding the master transcriptional regulator of flagellar synthesis (19–21). Accordingly, our BW25113-derived VHB17 strain, hereafter referred to as BW25113 *uspC*-*flhDC*::IS5, is motile under conditions where another BW25113 strain in our collection (VHB987) is largely non-motile (Fig. 1B). Genome sequencing of the non-motile BW25113 (VHB987) strain confirmed that it lacked the IS5 element upstream of the *flhDC* locus (Fig. 1A). Except for the lack of this insertion and a silent mutation in the *dnaE* gene, which is found only in the motile *uspC-flhDC*::IS5 strain, the two strains are otherwise isogenic (Table S1). Since some phages are flagellotropic, i.e., they initially attach to host flagella, we selected the motile BW25113 *uspC-flhDC*::IS5 strain as the host for phage isolation.

**Figure 1.**
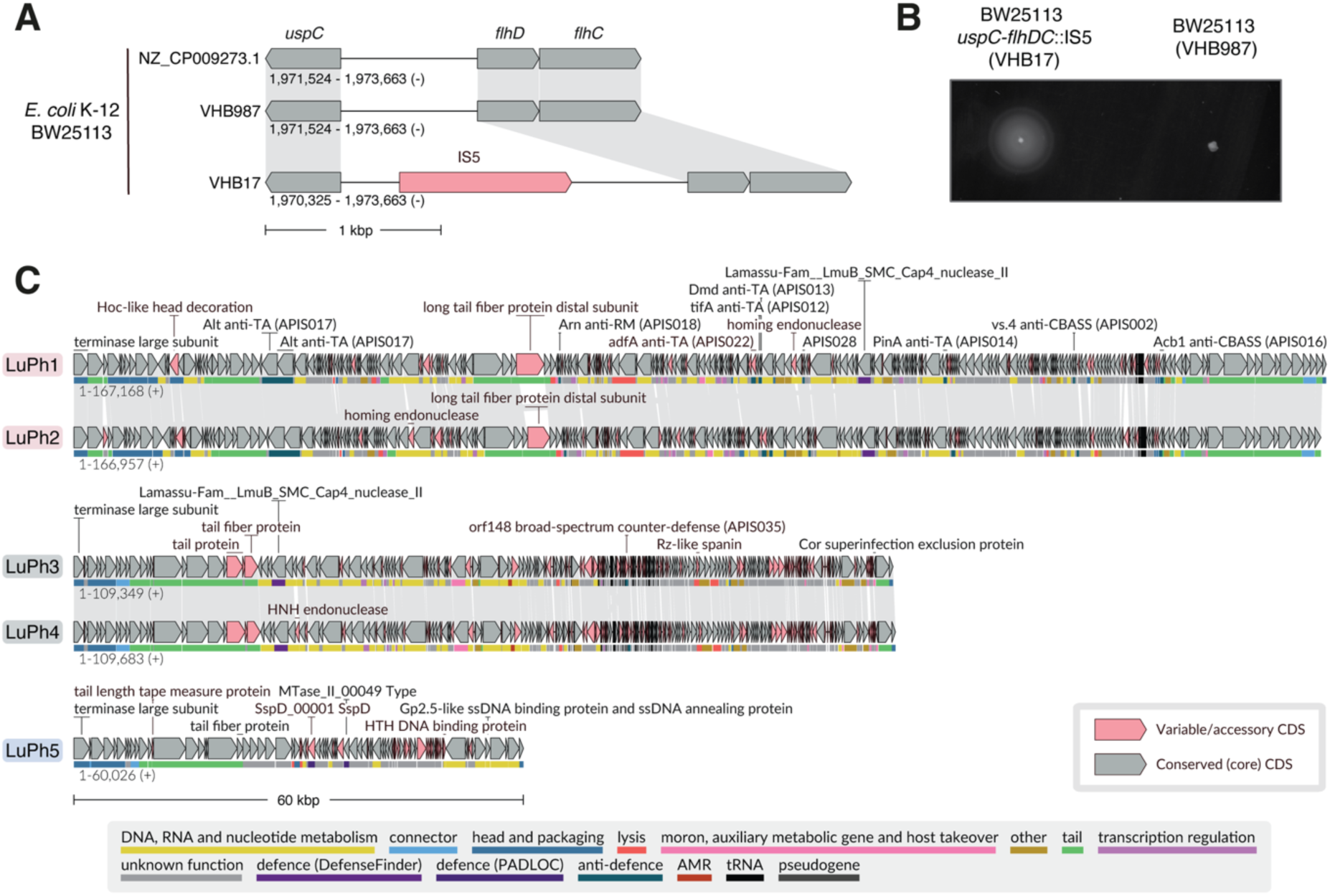
*flhDC* locus variation, motility phenotypes of *E. coli* BW25113 derivatives, and genome organization of the five LuPh phages. **(A)** Visualization of the *flhDC* locus in *E. coli* BW25113 derivatives VHB987 and VHB17, aligned to the reference BW25113 genome (NZ_CP009273.1). The IS5 mobile element is highlighted in coral pink. **(B)** Swimming motility assay on a soft agar plate. The plate was imaged after incubation at 37°C for 8 h. **(C)** Genome organization of LuPh phages. Conserved genes are shown in gray, and genes encoding variable protein groups are highlighted in coral pink. Functional annotations are indicated by colored lines beneath each ORF, corresponding to the color key at the bottom of the panel. A manually selected set of non-hypothetical encoded proteins are labeled, including the large terminase subunit, proteins identified as variable by LoVis4u, and proteins with functional hits based on PyHMMER annotation.

To isolate *E. coli*-infecting phages from the environment, water samples were collected from several small ponds in the Lund University Botanical Garden. These samples were cleared of debris, concentrated by zinc chloride precipitation, and assayed for phages using the top agar overlay method (22) using BW25113 *uspC*-*flhDC*::IS5 as the host. Individual phage isolates were purified by consecutive streaks of individual plaques. The genomes of 12 purified phages were sequenced using Illumina technology, then assembled and annotated. After removing duplicate isolates, five distinct *Escherichia* phages remained, which we named Lubas (LuPh1), Lucat (LuPh2), Lupin (LuPh3), Lucris (LuPh4), and Kompetensportalen (LuPh5) (Table 1, Fig. 1C). These five phages, hereafter referred to by their Lund Phage collection (LuPh) designations, were queried for the most similar phages in the NCBI nucleotide database. These searches identified various *Escherichia*, *Salmonella*, *Klebsiella*, and *Yersinia* phages with high similarity to the individual phages (Table S2-S6). Schematic comparisons of the LuPh and related phages are shown in Fig. S1-S3. To classify the phages, we used VIRIDIC (23) to determine the intergenomic similarities between LuPh isolates and their related phages (Fig. S4). According to the International Committee on Taxonomy of Viruses (ICTV) criteria (24), phages with an nucleotide identity, over their full genome length, higher than 70% are considered to belong to the same genus, while those with an identity above 95% are classified as the same species. Applying these criteria, LuPh1 and LuPh2 fall within the *Tequatrovirus* genus, LuPh3 and LuPh4 within the *Tequintavirus* genus, and LuPh5 within the *Chivirus* genus (Table 1). Moreover, LuPh1, LuPh3, and LuPh4 represent new species, with LuPh3 and LuPh4 belonging to the same species.

**Table 1.**
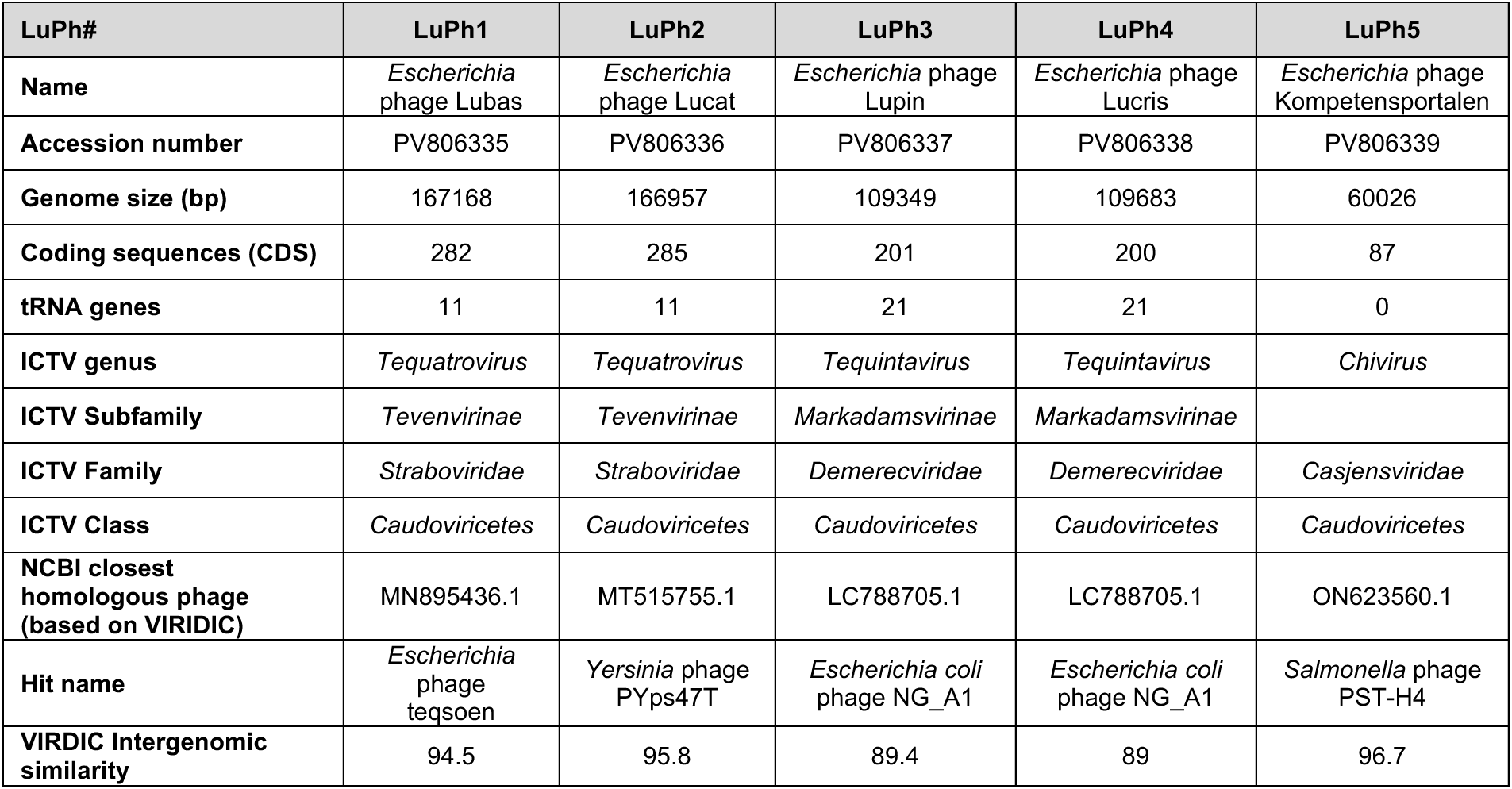
Phages isolated in this study.

Morphological characterization of the phages by transmission electron microscopy (TEM) showed that LuPh1 and LuPh2, consistent with their classification as *Tequatrovirus*, have long contractile tails (myoviruses) and prolate heads (Fig. 2A). The length of the tails and size of the heads are comparable between LuPh1 and LuPh2, with the latter observation aligning with their similarly sized genomes (Fig. 2B). As expected from their genus-level classification, LuPh3, LuPh4, and LuPh5 all possess long, flexible, non-contractile tails (siphoviruses) and icosahedral heads (Fig. 2A). Moreover, LuPh3 and LuPh4 exhibit the anticipated similarity in tail lengths and head sizes (Fig. 2B).

**Figure 2.**
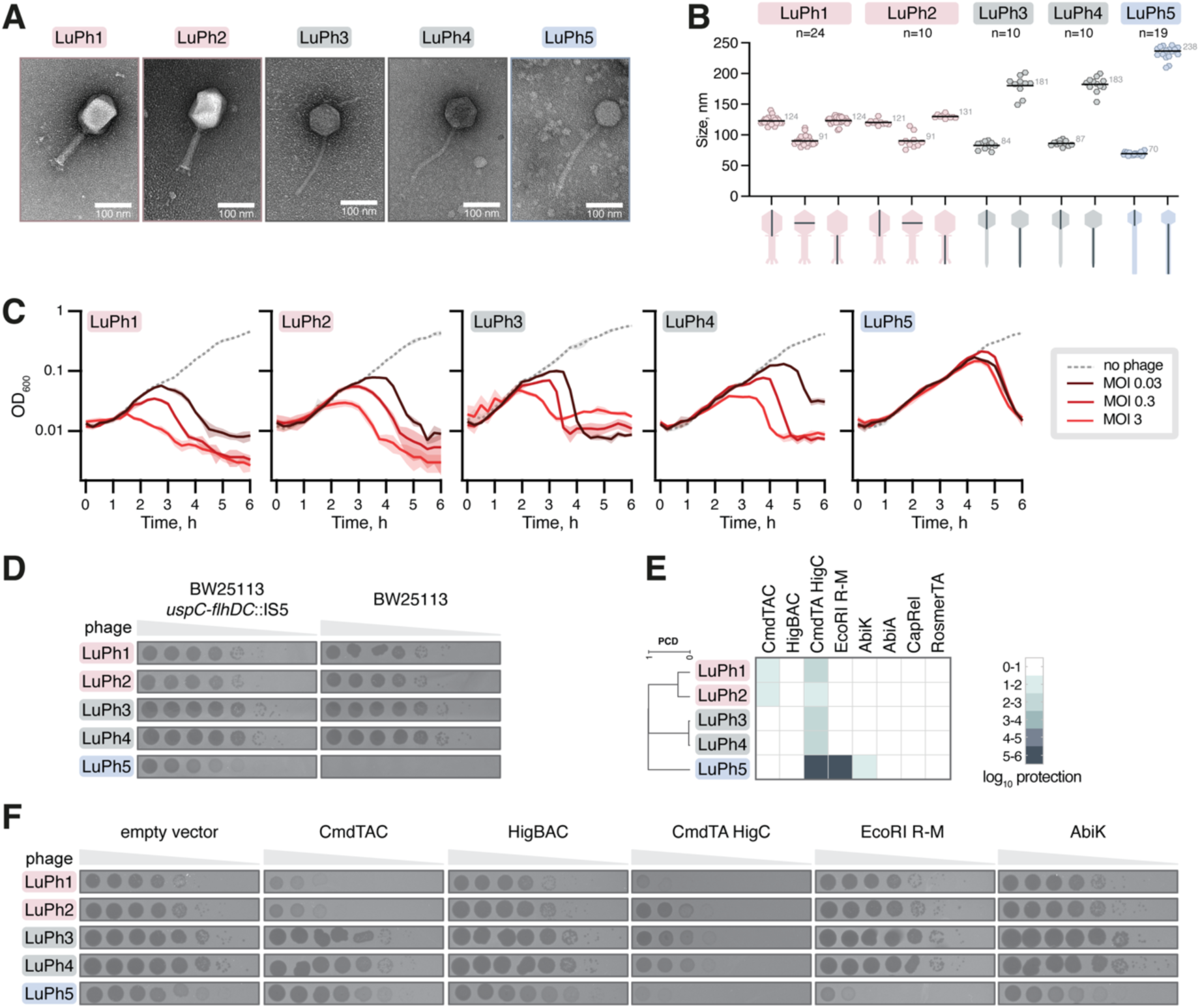
Morphology, infectivity, and anti-phage defense system-sensitivity of LuPh phages. **(A)** Representative transmission electron microscopy micrographs of LuPh phages **(B)** Dimensions of phage particles. Measured structural features are indicated below the x-axis. The mean value, from indicated individual measurements, is represented by a horizontal line and labeled in gray text. **(C)** Growth curves of *E. coli* BW25113 *uspC-flhDC*::IS5 (VHB17) in the presence of the indicated phages at MOIs of 0, 0.03, 0.3, and 3. Each curve represents the average of three biological replicates, each using a different transformant. Shaded areas indicate the standard deviation. **(D)** Serial dilution plaque assays on lawns of BW25113 *uspC*-*flhDC*::IS5 (VHB17) and BW25113 (VHB987). The LuPh phages were 10-fold serially diluted and spotted on the top agar plates followed by incubation at 37°C for 6 h. **(E, F)** Sensitivity of LuPh phages to selected anti-phage defense systems. *E. coli* BW25113 *uspC-flhDC*::IS5 (VHB17) harboring either an empty vector or a plasmid expressing the indicated anti-phage defense system were challenged with 10-fold serial dilutions of LuPh phages. Plates were imaged after incubation at 37°C for 6 h. Panel E shows a heatmap of log_10_ protection values, calculated as the mean of the log_10_- transformed efficiency of plating (EOP) values from three biological replicates, each using a different transformant. The dendrogram is defined by hierarchical clustering of the proteome composition distance (PCD) matrix. Panel (F) shows plaque assays from one of the replicates.

To assess the lytic activity of the LuPh phages, we performed liquid culture phage infection assays using multiplicities of infection (MOIs) ranging from 0.03 to 3. While all phages were capable of lysing BW25113 *uspC*-*flhDC*::IS5 cultures, LuPh5 showed a marked delay, with lysis occurring only during the late exponential phase (Fig. 2C). Although this delay may reflect a prolonged latency period of LuPh5, it could also be a consequence of the flagellotropic nature of bacteriophages in the *Chivirus* genus. As the number of flagella in *E. coli* increases during exponential growth and reaches a maximum in the late exponential phase (25), the delayed lysis could be a consequence of limited flagellar availability during early exponential phase. Since the LuPh phages were isolated on the motile BW25113 *uspC*-*flhDC*::IS5 strain, we also tested their ability to infect the largely non-motile BW25113 (VHB987) strain, which lacks the IS5 element between *uspC* and *flhDC*. The LuPh1, LuPh2, LuPh3, and LuPh4 phages showed comparable plaquing on both BW25113 derivatives (Fig. 2D). In contrast, LuPh5 failed to form plaques on the non-motile BW25113 strain (Fig. 2D; Supplementary Fig. S5 shows the same plaque assays after 24 h incubation at 37°C). The inability of LuPh5 to form plaques on the poorly motile BW25113 is reminiscent (and mechanistically analogous) of the inability of the *Chi* phage to infect non-flagellated *Salmonella* and *E. coli* strains (26–28)

To further investigate the features of the LuPh phages, we tested their sensitivity to a panel of eight anti-phage defense systems available within our laboratory. Specifically, we tested the *E. coli* toxin-antitoxin-chaperone (TAC) systems CmdTAC and HigBAC as well as the artificial hybrid CmdTA-HigC TAC system (29), the *E. coli* type II R-M system EcoRI (30), the *Lactococcal* abortive infection (Abi) reverse transcriptases AbiA and AbiK (31), the fused toxin-antitoxin defense system toxSAS CapRel^SJ46^ (32), and the RosmerTA of the *Gordonia*-infecting phage Kita (33). The anti-phage defense assays, using BW25113 *uspC*-*flhDC*::IS5 as the host, revealed that the LuPh phages showed sensitivity to a subset of the systems (Fig 2E and 2F; Supplementary Fig. S5 shows the same plaque assays as in Fig. 2F after 24 h incubation at 37°C). The CmdTAC system confers protection against LuPh1 and LuPh2, which is consistent with the previous finding that the system protects against several phages within the *Tequatrovirus* genus (29). All five phages are sensitive to the artificial CmdTA-HigC hybrid system, which is known to have a broader anti-phage defense spectrum than the parental systems (29). Finally, AbiK provides weak and EcoRI strong protection against LuPh5. While the mechanism by which AbiK protects against phages is unknown, the endonuclease of the EcoRI R-M system cleaves unmodified DNA at 5’-GAATTC-3’ sequences. LuPh5 harbors 25 EcoRI recognition sites in its genome and the sensitivity of the phage indicates that it does not harbor EcoRI-inhibiting base modifications or anti-defense systems.

In conclusion, we have characterized five environmental coliphages isolated during the 2023 *Fundamentals of Basic and Applied Phage Biology* course. These phages can potentially expand the toolkit for probing phage-host interactions. In particular, the use of the motile BW25113 *uspC*-*flhDC*::IS5 strain as the host enabled the isolation of LuPh5, a member of the *Chivirus* genus in the *Casjensviridae* family. As phages from the *Casjensviridae* family are absent in both the BASEL and Durham collections (9, 10, 13), LuPh5 could be a useful addition to the set of phages used to define the activity of candidate anti-phage defense systems. However, the inability of LuPh5 to infect the poorly motile BW25113 strain, lacking the IS5 element between *uspC* and *flhDC*, suggests that the host strain needs to be motile. Further work is needed to determine the host range for LuPh5 and the other LuPh phages.

## Materials and Methods

### Bacterial strains, plasmids, and media

The two *E. coli* strains used in this study (VHB17 and VHB987) represent differently sourced BW25113 (F– *λ^−^ Δ(araD-araB)567 Δ(rhaD-rhaB)568 ΔlacZ4787(::rrnB-3) hsdR514 rph-1*) strains. The differences compared to the reference BW25113 genome(16) are summarized in Table S1. Plasmids used in this study are listed in Table S7. The plasmid-borne CmdTAC, HigBAC, CmdTA-HigC, AbiK, AbiA, and RosmerTA systems are all under control of the constitutive P*_tet_* promoter whereas the EcoRI R-M system is expressed from the native promoter. The pJD1423_SD_CapRel^SJ46^ plasmid was constructed using Gibson assembly by cloning the CapRel^SJ46^-encoding ORF (32) under the control of the constitutive P*_tet_* promoter in pJD1423 (33). In pJD1423_SD_CapRel^SJ46^, the CapRel^SJ46^ ORF is preceded by the sequence for a strong Shine-Dalgarno element (5’-AGGAGGAATTAA-3’). *E. coli* strains were grown at 37°C in liquid or on solid (1.5% w/v agar) LB medium (10 g/L tryptone, 5 g/L yeast extract, and 10 g/L NaCl). When required for plasmid selection, the medium was supplemented with 100 μg/mL ampicillin.

### Bacterial motility assays

Overnight cultures of *E. coli* BW25113 *uspC-flhDC*::IS5 (VHB17) and BW25113 (VHB987), grown in LB medium, were diluted to an OD_600_ of 0.5. Subsequently, 1 μL of each culture was inoculated into a soft agar (LB solidified with 0.3% (w/v) agar) plate by stabbing the agar with a pipette tip and dispensing the culture while withdrawing the tip. Plates were incubated at 37°C for 8 hours.

### Bacterial genome sequencing

Genomic DNA from the VHB17 and VHB987 strains was purified using the NucleoSpin Microbial DNA Mini kit (Macherey-Nagel). Illumina sequencing was performed by SeqCenter (Pittsburgh, PA, USA), and Oxford Nanopore sequencing was conducted by Seqvision (Vilnius, Lithuania). Briefly, Illumina sequencing libraries were prepared using the tagmentation- and PCR-based Illumina DNA Prep kit and custom IDT 10bp unique dual indices (UDI) with a target insert size of 280 bp. No additional DNA fragmentation or size selection steps were performed. Sequencing was performed on an Illumina NovaSeq X Plus sequencer in one or more multiplexed shared-flow-cell runs, generating 2x151bp paired-end reads. Demultiplexing, quality control and adapter trimming was performed with bcl-convert (v4.2.4). For Oxford Nanopore sequencing, genomic DNA was end-repaired with the addition of dA overhangs and ONT kit 14 native barcodes were ligated by T/A ligation. The samples were purified, pooled, and nanopore native adapters were then ligated (kit 14 chemistry). Samples were sequenced on R10.4.1 nanopore flow cells and basecalled using the super-accuracy v5.0.0 model on Dorado v0.9.5. Reads were demultiplexed using Dorado. The lowest quality 10% of reads were removed using Filtlong v0.2.1.

Quality control of llumina paired-end and Nanopore reads was reanalyzed using FastQC v0.11.8 (34). Bacterial Nanopore sequencing reads were assembled using Flye v2.9.4 (35). Pilon v1.24 (36) was used to verify the Nanopore assemblies, with Illumina reads aligned to the assemblies using Bowtie2 v2.4.1 (37). Prokka v1.14.6 (38) was used to annotate the assembled genomes. To compare the sequenced bacterial genomes to the reference strain *E. coli* K-12 BW25113 (NZ_CP009273.1) and to each other, genomes were first aligned using minimap2 v2.29 (39), and genomic differences were subsequently annotated using SyRI v1.7.1 (40). Loci with differences were visualized using LoVis4u (41).

### Phage isolation and preparation of phage stocks

Environmental water samples (50 mL each) were collected from several small ponds in the Lund University Botanical Garden on April 24, 2023. Samples were cleared from debris by centrifugation at 8,000 × g for 5 minutes at room temperature. Phages in the supernatant were precipitated by adding 1 mL of 2 M ZnCl_2_, followed by incubation at 37°C for 30 minutes. Following centrifugation at 8,000 × g for 10 minutes at room temperature, the pellet was resuspended in 0.5-1 mL of SM buffer (0.1 M NaCl, 0.01 M MgSO_4_, 0.05 M Tris-HCl, pH 7.5) and agitated at 600 rpm for 15 minutes at 37°C in a thermoblock. The suspension was then assayed for phages using the top agar overlay method (22). Briefly, 200 μL of the suspension was mixed with 200 μL of an overnight culture of *E. coli* BW25113 *uspC-flhDC*::IS5 and incubated at room temperature for 15 minutes. The phage-cell mixture was then combined with 10 mL of top agar (0.5% w/v agar supplemented with 20 mM MgSO_4_ and 5 mM CaCl_2_) and poured onto square (12 × 12 cm) LB agar plates (1.5% w/v agar). Once the top agar had solidified, plates were incubated overnight at 37°C. Due to time constraints, individual plaques were re-streaked only once (using top agar overlays) during the course, but were subsequently purified by at least three consecutive re-streaks.

Phage stocks were prepared by transferring the top agar from visible lysis zones of the final re-streak into 1 mL of SM buffer in a 2 mL tube. The mixture was vortexed for 30 seconds and then centrifuged at 14,000 rpm for 10 minutes at room temperature. The resulting supernatant (i.e., the phage stock) was transferred to a 1.5 mL tube and stored at 4 °C. For long-term storage, the phages were kept as virocells at −80°C (42).

### Phage genome sequencing, annotation, and visualization

Genomic DNA from bacteriophages was purified using the Phage DNA Isolation Kit (Norgen Biotek). Illumina sequencing was conducted by SeqCenter (Pittsburgh, PA, USA). The sequencing libraries were prepared as described for bacterial genome sequencing above with the difference that the target insert size was 320 bp and that sequencing was performed on an Illumina NovaSeq 6000 system. Demultiplexing, quality control and adapter trimming was performed with bcl-convert (v4.1.5).

Additional quality control of paired-end reads was conducted using FastQC v0.11.8 (34). Reads were then trimmed for quality (-q 20) and filtered by length (--minimum-length 20) using cutadapt v2.10 (43). *De novo* genome assemblies were generated using SPAdes v3.15.2 (44) with default parameters, taking pre-processed paired-end reads as input. Genome assembly quality was assessed manually for each phage using Bandage v0.8.1 (45). Only single, circular, unbranched paths with predominant depths (∼1–2 × 10³) were retained as genome assemblies, while low-coverage, fragmented contigs were discarded.

All phages were annotated with Pharokka v1.5.1 (46). Specifically, coding sequences (CDS) were predicted with PHANOTATE (47), tRNAs were predicted with tRNAscan-SE 2.0 (48), and tmRNAs were predicted with Aragorn (49). Functional annotation was generated by matching each CDS to the PHROGs (50), VFDB (51), and CARD (52) databases using MMseqs2 (53) and PyHMMER (54, 55). Blastn (56) was used to query each phage genome against the ncbi nucleotide collection. Tables listing the most similar phages, ranked by total score, are provided in Table S2-S6.

The Lund Collection phages, along with a subset of the most closely related phages, were analyzed with VIRIDIC (23) to define intergenomic similarities with species and genus clusters. Genomic maps of the LuPh and closely related phages were visualized using LoVis4u (41). For additional functional annotation of proteins, LoVis4u searched with PyHMMER (55) against DefenseFinder (57), PADLOC (58), dbAPIS (59), and AMRFinder (60) databases. LoVis4u was also used to calculate proteome composition similarities among the Lund phages.

### Transmission electron microscopy

For negative staining, 4 µL of a filtered phage stock, appropriately diluted in SM buffer, was applied onto a carbon-coated 400-mesh copper grid (EMS) that had been glow-discharged in air for 60 seconds. The grids were blotted from the edge using Whatman grade 1 filter paper and negatively stained with 2% uranyl acetate for 60 seconds. The grids were imaged using a Thermo Scientific™ Talos™ L120C transmission electron microscope operating at 120 kV, which was equipped with a 4k × 4k Ceta CMOS camera. The dimensions of phage particles were measured using ImageJ (61).

### Liquid culture phage infection assays

Three independent overnight cultures of *E. coli* BW25113 *uspC*-*flhDC*::IS5 in LB medium supplemented with 10 mM MgSO_4_ and 2.5 mM CaCl_2_ were diluted to OD_600_≈0.075 in the same medium and 100 μl were transferred to wells of a 96-well plate. Phage stocks were diluted in SM buffer to generate final MOI values of 3, 0.3, and 0.03 when 10 μl of the dilution was added to a well. Ten μl of SM buffer were added to the wells where no phage was added. Bacterial growth was monitored at 37°C with double-orbital shaking at 400 rpm in a SPECTROstar Nano plate reader (BMG LABTECH) by measuring OD_600_ every 15 minutes for 6 hours.

### Anti-phage defense assays

Overnight cultures of *E. coli* BW25113 *uspC*-*flhDC*::IS5 (VHB17), carrying either the empty vector pBR322_ΔPtet (30), the empty vector VHp1423 (33), or the corresponding plasmid with an anti-phage system of interest, were mixed with top agar to a final concentration of 0.075 OD_600_ units/mL and overlaid onto LB agar plates (1.5% agar). Individual phage stocks were 10-fold serially diluted in SM buffer and 2.5 μL of each of the eight dilutions was spotted onto the solidified top agar. The experiments were performed in triplicate using different transformants for each replicate. Plates were incubated at 37°C, and plaque formation was monitored at 6 and 24 hours. Plaques were counted at both time points, and the efficiency of plating (EOP) was determined by dividing the plaque-forming units (PFU) for the system by those for the appropriate vector control. EOP values were transformed to a –log_10_ scale, and the log_10_ protection value in the heat map is the mean of the three repeats. Since the identity of the empty vector did not influence the results, the plaque assays shown in Fig. 2F and Fig. S5 include only one of the two empty vectors (pBR322_ΔPtet).

## Data availability

Genome sequences of LuPh1–LuPh5 and the *E. coli* BW25113 derivatives VHB17 and VHB987 have been submitted to GenBank and will be publicly available upon release under accession numbers PV806335, PV806336, PV806337, PV806338, PV806339, CP195190, and CP195189, respectively.

## Acknowledgements

We thank the Lund University Botanical Garden and its director Allison Perrigo for providing access to the ponds. The participants of the 2023 *Fundamentals of Basic and Applied Phage Biology* course are all acknowledged for their contributions. We thank NDPIA and specifically Debra Milton, Hanna Eriksson, Annasara Lenman and Louise Lindbäck as well as Klas Flärdh at the faculty of Biology, Lund University, for their support. The Cryo-EM for Life Sciences facility at Lund University and Lund University Bioimaging Centre (LBIC) are acknowledged for providing TEM support. This work was supported by the Carl Trygger Foundation (CTS24:3450 to M.J.O.J.), the Knut and Alice Wallenberg Foundation (2020-0037 to G.C.A. and V.H.), the Swedish Research Council (2022-01603, 2023-02353 and 2024-06071 to G.C.A.; 2021-01146 and 2024-06059 to V.H.), the Estonian Research Council (PRG2696 to V.H.), the Göran Gustafsson Foundation for Research in Natural Sciences and Medicine (the Göran Gustafsson Prize to V.H.), the Royal Physiographic Society of Lund (Endowments for the Natural Sciences, Medicine and Technology, number 45379 to A.A.E. and number 44198 to K.E.). NDPIA was funded by a grant from the Swedish Research Council (521-2013-8750). The computations were enabled by the Berzelius resource provided by the Knut and Alice Wallenberg Foundation at the National Supercomputer Centre, and by resources provided by the National Academic Infrastructure for Supercomputing in Sweden (NAISS), partially funded by the Swedish Research Council through grant agreement no. 2022-06725.

