## Supplemental Figure S2 for "Characterization of five environmental phages infecting *Escherichia coli* K-12 isolated during a phage biology training course"

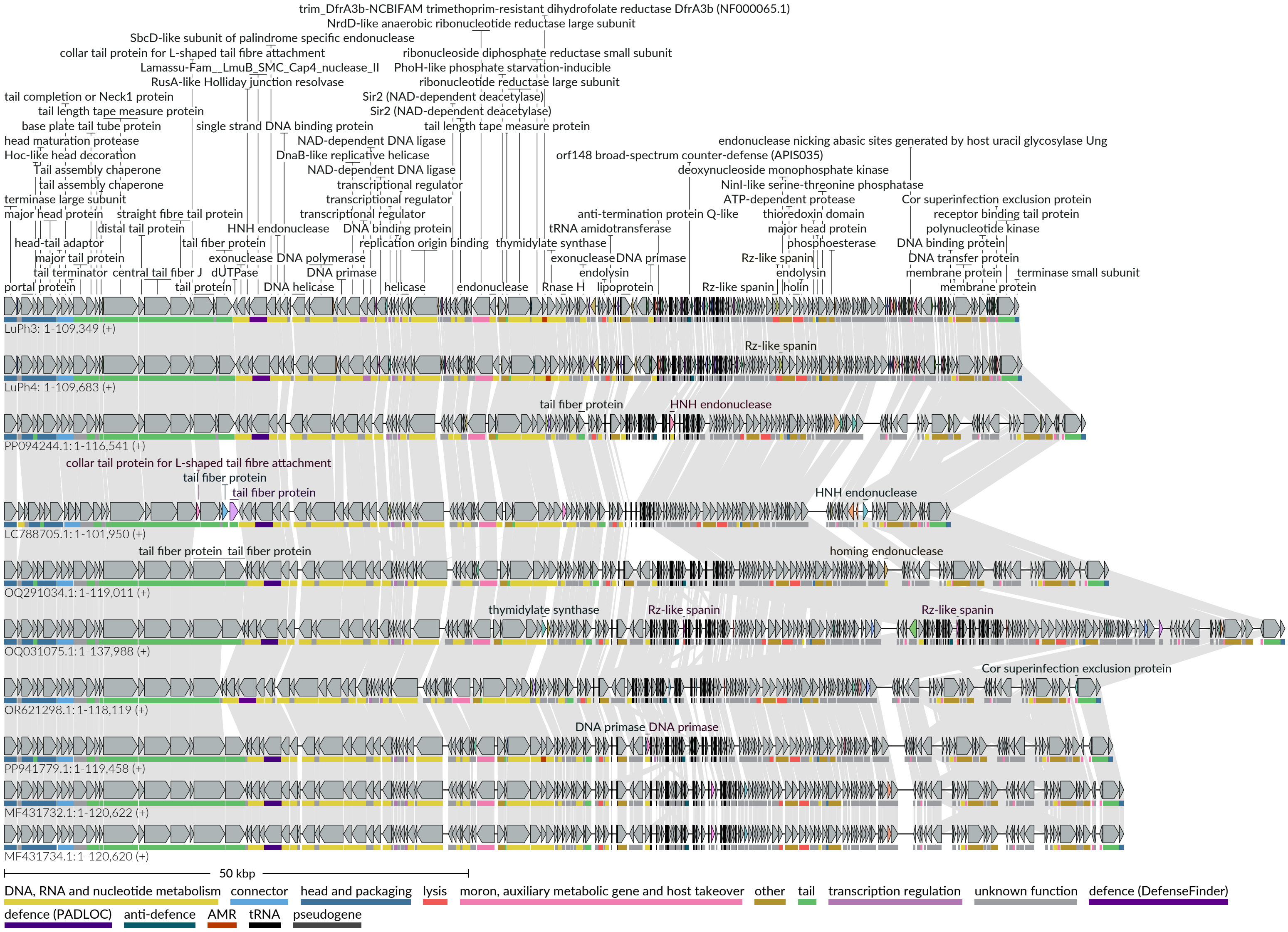


**Fig. S2.** Visualization of LuPh3 and LuPh4 and the eight most similar phages, generated using LoVis4u. Conserved genes are shown in gray, while genes encoding variable protein groups are highlighted with distinct colors. Protein classifications are derived from LoVis4u analysis of all LuPh3/4-like phages identified via BLAST search. Functional annotations are indicated by colored lines beneath each gene, following the color code at the bottom
