## Supplemental Figure S4 for "Characterization of five environmental phages infecting *Escherichia coli* K-12 isolated during a phage biology training course"

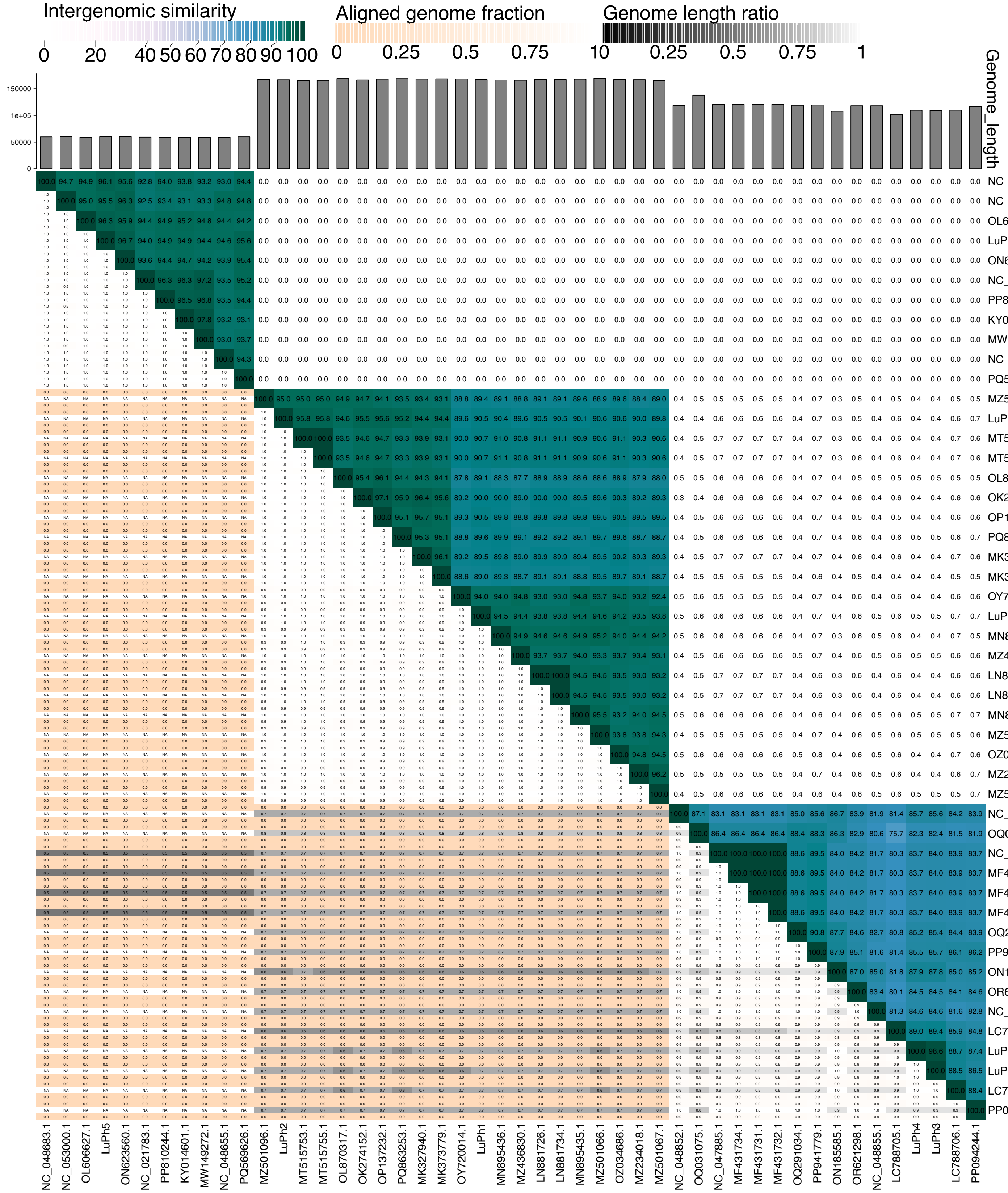


**Fig. S4.** Heatmap generated by VIRIDIC showing pairwise intergenomic similarities, aligned genome fractions, and genome lengths for a subset of the most similar phages to each LuPh group.
