## Supplemental Figure S5 for "Characterization of five environmental phages infecting *Escherichia coli* K-12 isolated during a phage biology training course"

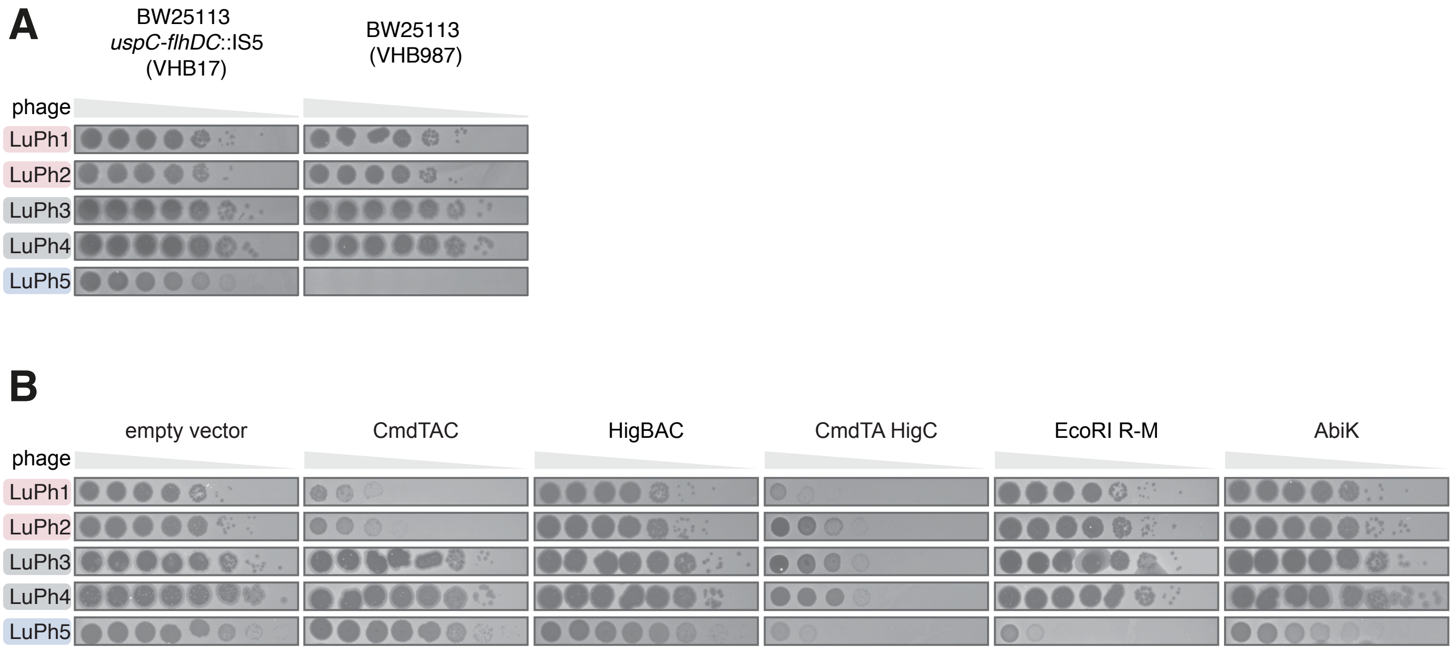


**Fig S5.** Serial dilution plaque assays. A) Plaque assays on lawns of BW25113 *uspC-flhDC*::IS5 (VHB17) and BW25113 (VHB987). The panel shows 24 h incubation of the of the assay shown in Fig. 2D. B) Plaques assays on lawns of BW25113 *uspC-flhDC*::IS5 (VHB17) harboring either an empty vector or a plasmid containing the indicated anti-phage defense system The panel shows 24 h incubation of the of the assay shown in Fig. 2F.
